# Spearfishing promotes a timidity syndrome and increases the safe operating distances in fish

**DOI:** 10.1101/205344

**Authors:** Valerio Sbragaglia, Lorenzo Morroni, Lorenzo Bramanti, Boris Weitzmann, Robert Arlinghaus, Ernesto Azzurro

## Abstract

In a landscape of fear, humans are altering key behaviors expressed by wild-living animals, including those related to foraging, reproduction and survival. When exposed to potentially lethal human actions, such as hunting or fishing, fish and wildlife is expected to behaviorally respond by becoming more timid, but proving such responses underwater in the wild has been challenging. Using a rich dataset collected *in situ*, we provide evidence of spearfishing-induced behavioral effects in five coastal fish species using the flight initiation distance (FID) as a proxy of predator avoidance and boldness. We document that spearfishing promotes a timidity syndrome (i.e., an increase of the average timidity of harvested populations) and that the wariness of prey’s wariness is influenced by individual size, level of protection offered through marine protected areas and the ability to recognize the risk posed by underwater human predators. In particular, we show that changes in the appearance of the observer (spearfisher *vs*. snorkeler) modulate the risk perception among the exploited species, and these differences are more evident outside marine protected areas where spearfishing is allowed. We also detected a positive correlation between FID and fish size, with larger specimens (that are more likely targets of spearfishers) revealing larger FID. The behavioral effects were most clearly expressed in the most heavily exploited species and declined towards the less desired and less targeted ones, which may be a result of learning mechanisms and plasticity and/or fisheries-induced evolution of timidity. Our study reveals a trade-off where intensive spearfishing negatively affects future spearfishing success through behavior-based alteration of catchability. Either rotating harvest or implementation of mosaics of protected and exploited areas might be needed to manage spearfishing-induced timidity in exploited stocks.

## INTRODUCTION

Human activities, such as hunting and fishing, are affecting wild populations by altering abundance, biomass, size-structure, population dynamics as well as life-histories and behavioral traits and associated demographic and evolutionary processes (Sullivan et al., 2017). A large body of literature has studied ecological (e.g. Worm et al., 2009, Arlinghaus et al., 2017) and evolutionary effects of fishing (e.g. Jørgensen et al., 2007, Laugen et al., 2014). Fishing is typically selective with respect to different traits such as size and behavior (Law, 2007, Kuparinen and Festa-Bianchet, 2017, Uusi-Heikkilä et al., 2008, Lennox et al., 2017), and elevated mortality as well as trait-selective harvesting is expected to affect size and age structure (e.g. Arlinghaus et al., 2010) as well as life-history and behavioral traits (Allendorf and Hard, 2009). For instance, many fishing methods selectively remove large and bold individuals (e.g. Klefoth et al., *in press*), altering traits of populations by favoring fast life-histories (Laugen et al., 2014, Heino et al., 2013) and more cautious behavioral types (Arlinghaus et al., 2017). These changes can be caused both by plastic responses as well as evolutionary changes (Laugen et al., 2014, Arlinghaus et al., 2017).

While fisheries-induced changes in life-histories have seen abundant attention in the literature, the possibility of fisheries altering the composition of fish behavioral types has been less explored (Heino et al., 2015). Recently, it been proposed that behavior is a sensitive indicator of exploitation in the marine environment (Goetze et al., 2017). The exploitation-induced behavioral effect has recently been defined as a "timidity syndrome", which may have a range of ecological, social and economic consequences, most notably by altering the catchability of exploited populations (Arlinghaus et al., 2017). However, despite considerable conceptual appeal, there are surprisingly few field studies that have provided clear documentation that exploited fish stocks are indeed more timid (Januchowski-Hartley et al., 2011, Alós et al., 2015, Alós et al., 2014, Tsuboi et al., 2016, Alós et al., 2016, Bergseth et al., 2016, Colefax et al., 2016), motivating the present work.

We investigated the behavioral changes induced by recreational spearfishing in several coastal sites in two Mediterranean countries. Spearfishing constitutes a globally popular recreational activity with important historical and social roots, especially in the Mediterranean Sea (Coll et al., 2004, Sbragaglia et al., 2016, Lloret et al., 2008). Spearfishing constitutes an active fishing mode sharing similarities to hunting because spearfishers are able to visually target and select their prey (Dalzell, 1996). The size-selectivity of spearfishing depends on the hunting technique (shelter seeking, ambush, sit-and-wait or direct approach) and is related to the fisher’s experience (Diogo et al., 2017). Similar to trophy hunting (e.g. Coltman et al., 2003), spearfishers select the large individuals, and as the capture depends on physical proximity of the fish and the spearfisher there is also the potential for selection on behavioral phenotypes, such as boldness, exploration or aggression (e.g. Januchowski-Hartley et al., 2011). Therefore, in heavily exploited areas, the traits linked to the ability of fishes to hide or escape from the human predator can be under a strong selective pressure (Ydenberg and Dill, 1986). Fish behavior has a large plastic component and can thus rapidly change through learning and development. As a consequence, levels of wariness can substantially increase due to increased spearfishing pressure within few days after opening a fishing zone (Goetze et al., 2017).

One of the most commonly used metric to assess a prey's wariness in the presence of (human or natural) predation, is the distance to which a predator can approach before the prey flees. This measure, indexed as the flight initiation distance (FID; Ydenberg and Dill, 1986), has been used in previous studies to estimate fish behavioral responses to spearfishers (Januchowski-Hartley et al., 2011, Januchowski-Hartley et al., 2012, Tran et al., 2016, Benevides et al., 2016, Feary et al., 2011, Gotanda et al., 2009, Nunes et al., 2015, Guidetti et al., 2008, Cole, 1994) and may constitute a sensitive indicator of exploitation to inform management (Januchowski-Hartley et al., 2014, Goetze et al., 2017). Generally, individuals with shorter FID are considered bolder (Januchowski-Hartley et al., 2011, Feary et al., 2011). Several studies have investigated how FID can be affected by natural and human factors, such as fishing pressure (key finding is an increased FID under fishing pressure), protection status of a given site (shorter FID in marine protected areas) and body size (increased FID with size) (Januchowski-Hartley et al., 2011, Januchowski-Hartley et al., 2012, Feary et al., 2011, Gotanda et al., 2009). Changes in the FID can be caused by a combination of both selection and evolution and plasticity within the realm of learned predator avoidance, predator recognition and inspection behaviors (Kelley and Magurran, 2003). No matter which mechanism, a spearfishing-induced increase in the FID has been repeatedly reported and will negatively affect future catchability and exposure of fishes to spearfishers (Feary et al., 2011).

The rapid increase of spearfishing in the last 60 years in the Mediterranean area (Coll et al., 2004) represents a new pressure for wild coastal fish populations but also an opportunity to understand how fish cope with the sudden arrival of a new predator, the underwater human predator. Fishes can became aware of the presence of predators by olfactory (Chivers and Smith, 1998), tactile or auditory stimuli (Kelley and Magurran, 2003), but in clear water conditions typical of many areas of the Mediterranean Sea, visual cues are likely the primary source of information that allows fishes to recognize predators from long distances (Aksnes and Giske, 1993). Fishes quickly learn how to reduce predation risk, thereby moderating the fundamental growth-mortality trade-off (Werner and Hall, 1974, Ydenberg and Dill, 1986, Stamps, 2007). Learning may be both an individual or social process, contributing to the large plasticity in behavioral repertoire that is typical of many fish species (Odling-Smee and Braithwaite, 2003, Kieffer and Colgan, 1992, Mangel and Stamps, 2001). However, a large body of literature has also revealed a high individual variability in behavioral traits, typically referred to as behavioral types or personalities (Carere and Maestripieri, 2013). Although personality can emerge from clonal fishes reared in apparently identical conditions (e.g. Bierbach et al., 2017), consistent variation in behavioral traits has been found to carry a large genetic (i.e., heritable) component (Dochtermann et al., 2015). Thus, spearfishing-induced selection on behavior could also induce evolutionary change towards timidity (Andersen et al., 2017).

We investigated the FID expressed across several species in the wild, in different geographical areas and under different levels of protection offered by marine protected areas where spearfishing is not allowed. To mechanistically understand how fishes respond to human-induced predation threat, we included a ‘spearfishing’ treatment, which allowed testing the ability of fishes to recognize the threat represented by humans. Our predictions were: (*i*) FID increases according to the threat level associated by fishes to the encounter with spearfishers (higher FID in the presence of a diver with a speargun; higher FID outside the marine protected areas); (*ii*) FID increases in proportion to the historic spearfishing pressure on a given species (the higher the historic exploitation on a given species, the higher is the FID); and (*iii*) FID increases with fish size because larger fishes are preferentially targeted by spearfishers (the larger the fish, the higher the FID).

## MATERIALS AND METHODS

### Study species

We targeted five common coastal fish taxa subjected to different levels of spearfishing harvesting in the following order (from low to high historic exploitation intensity): (1) *Symphodus tinca;* (2) *Sarpa salpa*; (3) mugilidae (considering the following species: *Chelon labrosus, Mugil cephalus, Liza aurata* and *Liza ramada*); (iv) *Diplodus vulgaris*; and (v) *Diplodus sargus.* This gradient was compiled on the basis of literature (Harmelin et al., 1995, Coll et al., 2004, Lloret et al., 2008, Rocklin et al., 2011) and reflecting the authors’ knowledge of spearfishers’ habits in the studied areas. FID was measured in a 2×2 factorial design in two areas with different level of harvesting protection status (protected/unprotected sites) and in two diver treatments (potentially threatening spearfishing and non-threatening snorkeling). These treatments and sites represented a gradient of threatening situations for the fishes: strong threat (unprotected/spearfishing: hereafter Spear-NP), medium threat (unprotected/snorkeling: hereafter Snork-NP), weak threat (protected/spearfishing: hereafter Spear-P) and very weak threat (protected/snorkeling: hereafter Snork-P).

### Study sites and experimental design

The FID of the five target species was surveyed between May and October 2016 in 3 different marine protected areas (MPAs) of the North-West Mediterranean Sea. The study was conducted in the MPA of Bonifacio Straits, Corsica, France (protected zone: 41° 35.28’ N, 9° 21.76’E; non-protected reference zone: 41° 36.99’N, 9° 21.19’ E), the MPA of Cerbère-Banyuls, France (protected zone: 42° 28.58 N, 3° 9.40’E; non-protected reference zone: 42° 29.31’N, 3°7.79’ E) and the MPA of Medes Islands, Spain (protected zone: 42° 2.67’N, 3° 13.40’E; non-protected zone: 42° 6.20°N, 3° 10.50’E). We chose three MPAs and two countries to generalize our findings and reduce the impact of specific uncontrolled contextual factors within any given site/country. Hereafter, we will refer to the three different areas as Banyuls, Bonifacio and Medes.

The FID was surveyed inside and outside each MPA. All observations were conducted at a minimum distance of 750 m from the MPA borders, to avoid potential spillover of naive fish from the MPA (Alós et al., 2015, Januchowski-Hartley et al., 2013). All sites were surveyed by the same operator (LM). The spearfisher was equipped with a speargun (100 cm long, rubbers were removed and substituted with PVC tubes to not violate regulations of the MPA) and with the typical spearfishing equipment (black wetsuit, long black fins and a little black mask). The snorkeler was equipped with a shorty bluish wetsuit, short bluish fins and a large colored mask. The different diver configurations (spearfisher/snorkeler) were applied inside and outside each MPA in a random order over 4-7 consecutive days.

### Data collection

The operator swam on the surface along a linear transect over rocky reef bottom between 1 and 3 m depth, identifying the target fish from the surface, and then swimming towards it at a steady speed (Januchowski-Hartley et al., 2014). When the fish reacted to the presence of the operator (usually flight away but in few cases, in the marine protected areas, positive interactions also occurred) the position was marked on the bottom with a small weight. Positive interactions were not considered in this study. The distance between the location of escape and the marked position was then measured with a tape and recorded as FID (Ydenberg and Dill, 1986). Flight was considered to have occurred when the fish increased its swimming speed or changed its swimming direction in response to the operator. For each target fish (<10 cm), the species was determined and total length was estimated to the nearest cm. To establish the accuracy of the size estimation, the length of known size static objects (length range 10-70 cm; n = 30) was estimated with an error of 8.4 ± 6.4% (average percent error ± standard deviation). Larger mobile species and individuals were counted first to minimize the chance of approaching a target fish that had been disturbed by a previous trial, consecutive trials were conducted a minimum of 10 m apart, and in the opposite direction to which a disturbed fish fled, assuming independence of samples within sites.

### Statistical analysis

We implemented a linear mixed model with random intercepts using FID as response variable, fish length as covariate, treatment (four levels: Spear-NP; Snork-NP; Spear-P; Snork-P) and taxa (five levels *D. sargus*, *D. vulgaris*, *Mugilidae*, *S. salpa* and *S. tinca*) as fixed effects, and the three areas (Bonifacio, Banyuls and Medes) as random effect. Model fitting was examined by checking normality of residuals and plotting theoretical quantiles versus standardized residuals. We used the “lme4” R package (Bates et al., 2014) to implement the linear mixed models, and “lsmeans” R package (Lenth, 2016) to run multicomparisons among slopes or means. All analyses were conducted in R 3.3.1 (R Core Team 2016) with a 95% confidence interval.

## RESULTS

The FID of N = 1,328 fishes (345 *S. tinca*, 177 *S. salpa*, 130 mugilidae, 286 *D. vulgaris* and 390 *D. sargus*) distributed among areas and treatments was scored (Table 1). All the fixed terms revealed significant effects on FID (Table 2). Most importantly, a significant three-way interaction effect among size, treatment and species was detected (χ_12,1328_ = 37.47, p < 0.0001; Table 2), indicating that the treatments exerted differential effects depending on the size of the fish and the species.

**Table 1.** The number of observations collected for each species or group of species in each Western Mediterranean areas (Banyuls, Bonifacio, Medes) and for each treatment (Snork-P: snorkeling inside the marine protected area; Spear-P: spearfishing inside the marine protected area; Snork-NP: snorkeling outside the marine protected area; Spear-NP: spearfishing outside the marine protected area)

| Species | Banyuls |  |  |  | Bonifacio |  |  |  | Medes |  |  |  | Total |
| --- | --- | --- | --- | --- | --- | --- | --- | --- | --- | --- | --- | --- | --- |
|  | Snork-P | Spear-P | Snork-NP | Spear-NP | Snork-P | Spear-P | Snork-NP | Spear-NP | Snork-P | Spear-P | Snork-NP | Spear-NP |  |
| *S. tinca* | 29 | 28 | 31 | 30 | 43 | 29 | 28 | 26 | 27 | 25 | 19 | 30 | 345 |
| *S. salpa* | 19 | 8 | 7 | 14 | 21 | 18 | 11 | 9 | 23 | 17 | 13 | 17 | 177 |
| *mugilidae* | 18 | 17 | 3 | 9 | 3 | 5 | 8 | 11 | 22 | 20 | 5 | 9 | 130 |
| *D. vulgaris* | 34 | 19 | 30 | 47 | 20 | 20 | 15 | 19 | 21 | 13 | 20 | 28 | 286 |
| *D. sargus* | 68 | 37 | 32 | 37 | 34 | 30 | 20 | 16 | 25 | 33 | 32 | 26 | 390 |

**Table 2.** The results of the liner mixed model implemented for Flight Initiation Distance (FID). The fixed factors, the covariate and their interactions are reported together with degree of freedom (df), the Chi-square value statistics (X^2^) and the *p* value (*p*). The overall mean estimation of FID is reported for each species together with standard error in parentheses. Letters indicate the output of the Tukey post hoc test (a > b > c). See Table 3 for further results.

| Predictor variables | df | $X^2$ | $p$ |
| --- | --- | --- | --- |
| Size | 1 | 390.19 | < 0.0001 |
| Species | 4 | 379.32 | < 0.0001 |
| Treatment | 3 | 858.54 | < 0.0001 |
| Size × Treatment | 3 | 52.96 | < 0.0001 |
| Size × Species | 4 | 66.10 | < 0.0001 |
| Treatment × Species | 12 | 115.27 | < 0.0001 |
| Size × Treatment × Species | 12 | 37.47 | < 0.0001 |

The mean FID, while accounting for the mean effect of size, increased outside the MPA both in the presence of the spearfisher and snorkeler for the historically most targeted species (*D. sargus, D. vulgaris* and *S. salpa*; Table 3, Figs. 2, 3a), while for the historically less targeted mugilidae species and *S. tinca* the FID increase was recorded only in the presence of the spearfisher (Table 3, Figs. 2, 3a). Outside the MPA, the mean FIDs were always larger than 4 m (except for *S. tinca*; see Table 3). Inside the MPA, however, the FID was not significantly different in the presence of either spearfisher or a non-threatening snorkeler, with the exception of the reaction of *S. tinca* (Table 1, Figs. 2, 3a). By contrast, the presence of a spearfisher outside the MPA increased the mean FID in all the species except for the less targeted *S. salpa* and *S. tinca* (Table 3, Figs. 2, 3a).

**Table 3.**
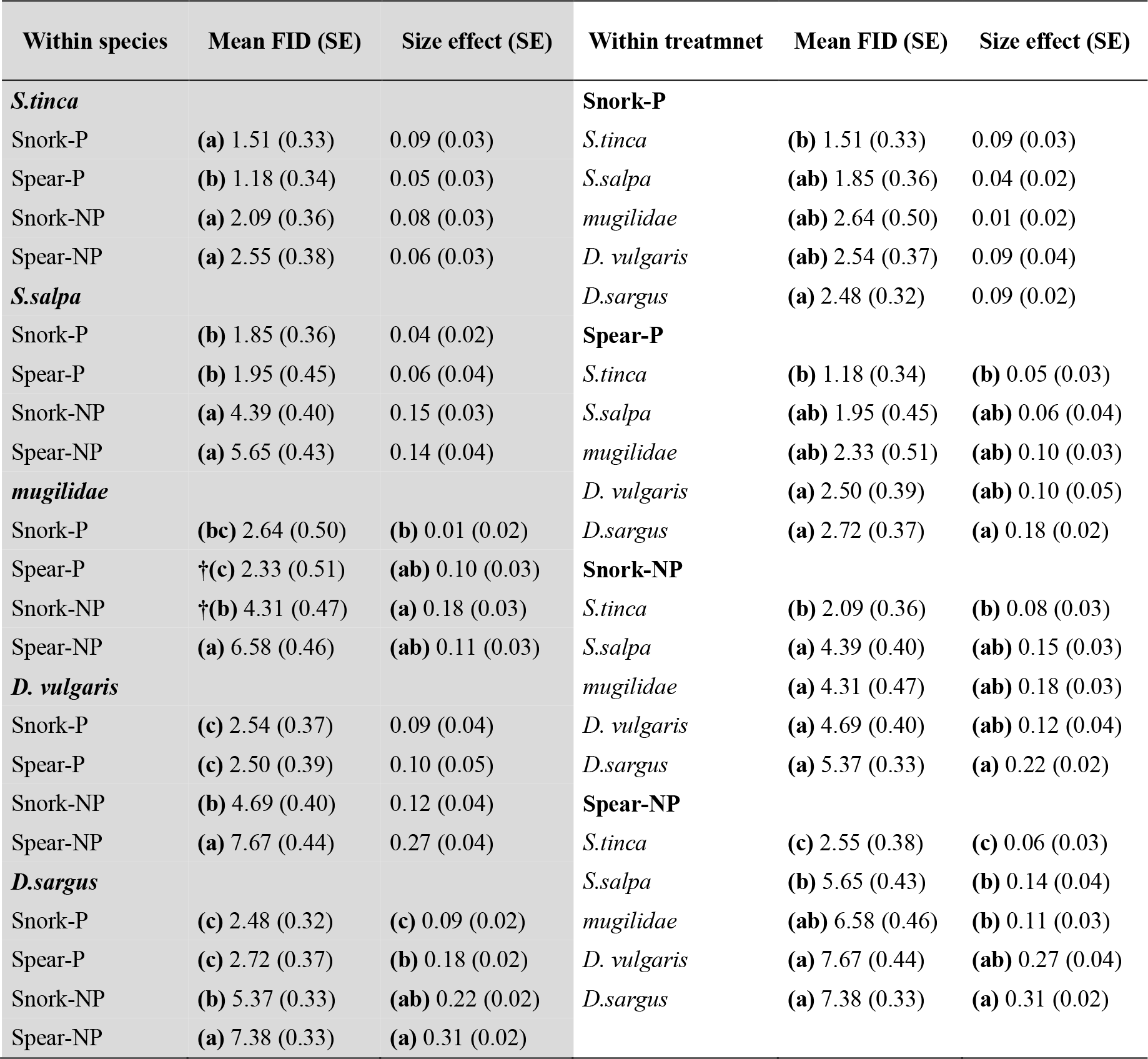
The results of the linear mixed model implemented for flight initiation distance (FID). The estimations of FID are related to the significant three way interactions reported in Table 2 (Size × Treatment × Species). The estimation of FID within the same species along the treatments are reported in the left part of the table (grey area), while the estimation of FID within the same treatment along the species are reported in the right part (white area). Species are listed according to an increasing gradient of spearfishing pressure and treatments (Snork-P: snorkeling inside the marine protected area; Spear-P: spearfishing inside the marine protected area; Snork-NP: snorkeling outside the marine protected area; Spear-NP: spearfishing outside the marine protected area) following an increasing gradient of threat. The mean FID values adjusted for the average size (22.5 cm) and the effect of size on FID (slope) are reported together with the standard error in parentheses. Letters indicate the output of the Tukey post hoc test (a > b > c). † Indicates a trend between groups (p<0.10).

| Within species | Mean FID (SE) | Size effect (SE) | Within treatment | Mean FID (SE) | Size effect (SE) |
| --- | --- | --- | --- | --- | --- |
| ***S.tinca*** |  |  | **Snork-P** |  |  |
| Snork-P | (a) 1.51 (0.33) | 0.09 (0.03) | *S.tinca* | (b) 1.51 (0.33) | 0.09 (0.03) |
| Spear-P | (b) 1.18 (0.34) | 0.05 (0.03) | *S.salpa* | (ab) 1.85 (0.36) | 0.04 (0.02) |
| Snork-NP | (a) 2.09 (0.36) | 0.08 (0.03) | *mugilidae* | (ab) 2.64 (0.50) | 0.01 (0.02) |
| Spear-NP | (a) 2.55 (0.38) | 0.06 (0.03) | *D. vulgaris* | (ab) 2.54 (0.37) | 0.09 (0.04) |
| ***S.salpa*** |  |  | *D.sargus* | (a) 2.48 (0.32) | 0.09 (0.02) |
| Snork-P | (b) 1.85 (0.36) | 0.04 (0.02) | **Spear-P** |  |  |
| Spear-P | (b) 1.95 (0.45) | 0.06 (0.04) | *S.tinca* | (b) 1.18 (0.34) | (b) 0.05 (0.03) |
| Snork-NP | (a) 4.39 (0.40) | 0.15 (0.03) | *S.salpa* | (ab) 1.95 (0.45) | (ab) 0.06 (0.04) |
| Spear-NP | (a) 5.65 (0.43) | 0.14 (0.04) | *mugilidae* | (ab) 2.33 (0.51) | (ab) 0.10 (0.03) |
| ***mugilidae*** |  |  | *D. vulgaris* | (a) 2.50 (0.39) | (ab) 0.10 (0.05) |
| Snork-P | (bc) 2.64 (0.50) | (b) 0.01 (0.02) | *D.sargus* | (a) 2.72 (0.37) | (a) 0.18 (0.02) |
| Spear-P | †(c) 2.33 (0.51) | (ab) 0.10 (0.03) | **Snork-NP** |  |  |
| Snork-NP | †(b) 4.31 (0.47) | (a) 0.18 (0.03) | *S.tinca* | (b) 2.09 (0.36) | (b) 0.08 (0.03) |
| Spear-NP | (a) 6.58 (0.46) | (ab) 0.11 (0.03) | *S.salpa* | (a) 4.39 (0.40) | (ab) 0.15 (0.03) |
| ***D. vulgaris*** |  |  | *mugilidae* | (a) 4.31 (0.47) | (ab) 0.18 (0.03) |
| Snork-P | (c) 2.54 (0.37) | 0.09 (0.04) | *D. vulgaris* | (a) 4.69 (0.40) | (ab) 0.12 (0.04) |
| Spear-P | (c) 2.50 (0.39) | 0.10 (0.05) | *D.sargus* | (a) 5.37 (0.33) | (a) 0.22 (0.02) |
| Snork-NP | (b) 4.69 (0.40) | 0.12 (0.04) | **Spear-NP** |  |  |
| Spear-NP | (a) 7.67 (0.44) | 0.27 (0.04) | *S.tinca* | (c) 2.55 (0.38) | (c) 0.06 (0.03) |
| ***D.sargus*** |  |  | *S.salpa* | (b) 5.65 (0.43) | (b) 0.14 (0.04) |
| Snork-P | (c) 2.48 (0.32) | (c) 0.09 (0.02) | *mugilidae* | (ab) 6.58 (0.46) | (b) 0.11 (0.03) |
| Spear-P | (c) 2.72 (0.37) | (b) 0.18 (0.02) | *D. vulgaris* | (a) 7.67 (0.44) | (ab) 0.27 (0.04) |
| Snork-NP | (b) 5.37 (0.33) | (ab) 0.22 (0.02) | *D.sargus* | (a) 7.38 (0.33) | (a) 0.31 (0.02) |
| Spear-NP | (a) 7.38 (0.33) | (a) 0.31 (0.02) |  |  |  |

**Figure 1.**
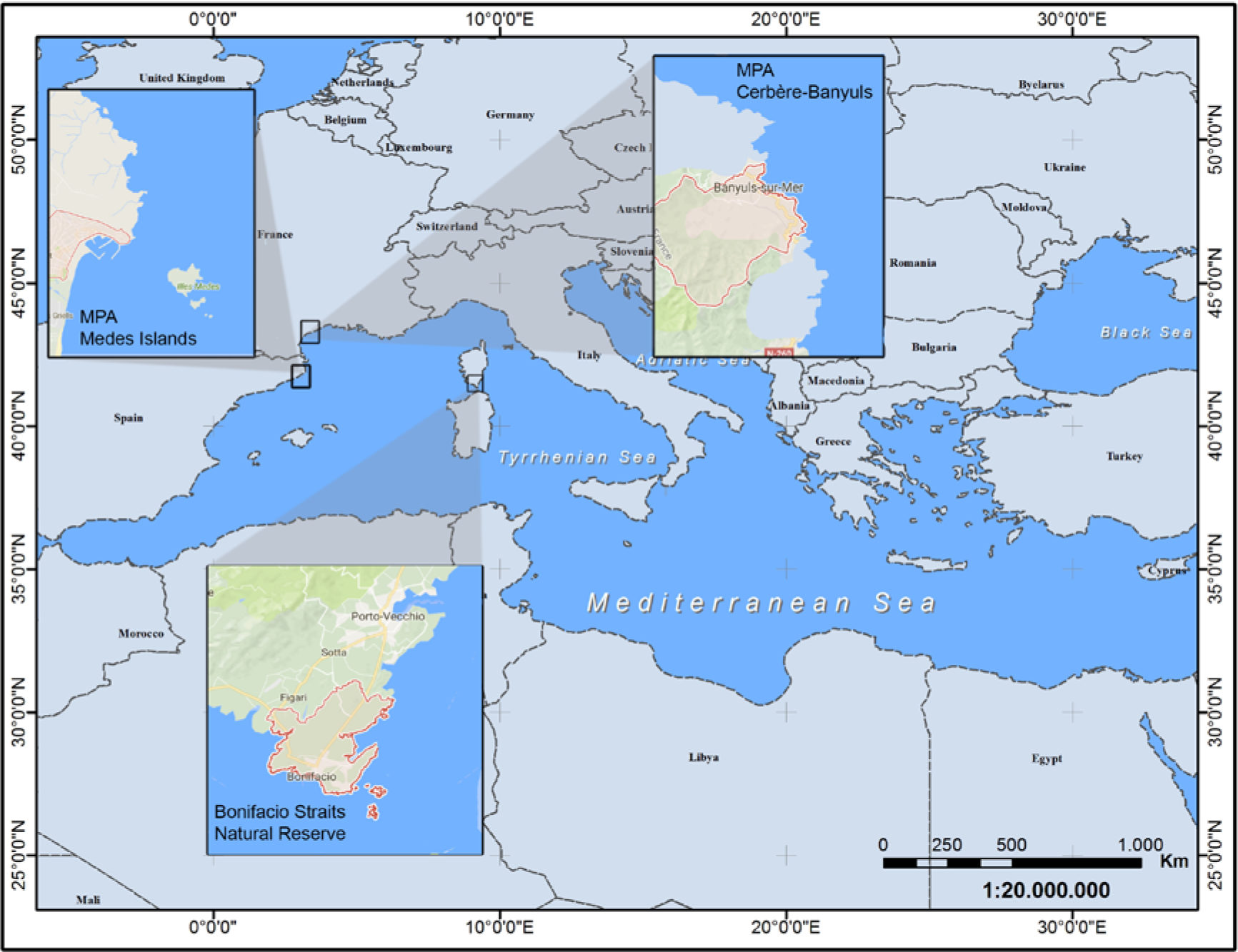
Areas of study. Surveys were performed in and out three protected areas: the Bonifacio Straits Natural Reserve (Corsica), the Marine Protected Area of of Cerbère-Banyuls (France) and the Marine Protected Area of Medes islands (Spain) in the Mediterranean Sea.

**Figure 2.**
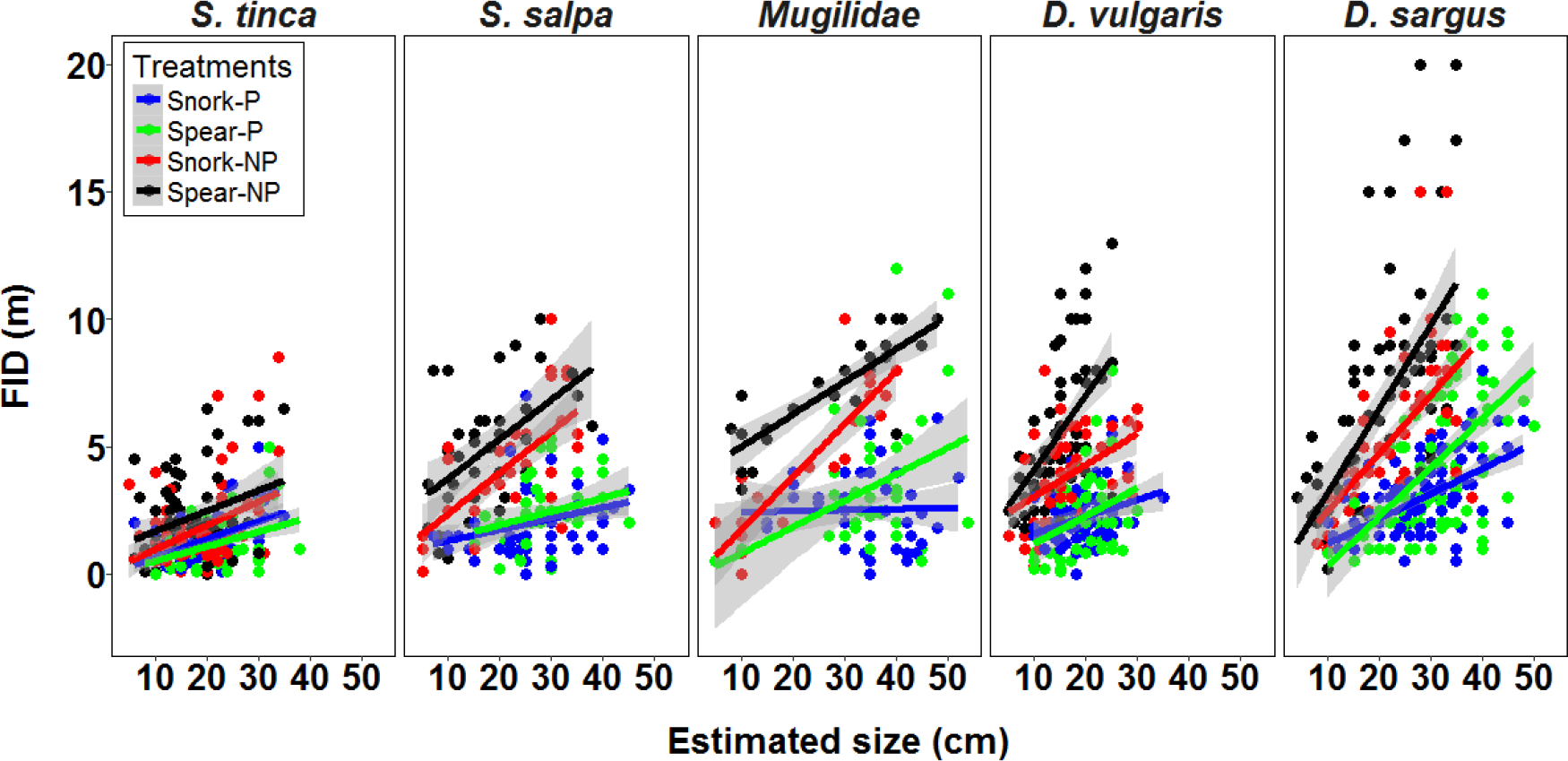
The raw flight initiation distance measurements (dots) are presented for each species listed according to an increasing gradient of spearfishing pressure for the four treatments in function of the estimated size of fishes. The different colors represent the four treatments (blue: snorkeling in marine protected area; green: spearfishing in marine protected area; red: snorkeling outside marine protected area; black: spearfishing outside marine protected area) used in this study. The solid lines represent the linear correlation between FID and size expressed together with the 95% confidence interval (grey area).

**Figure 3.**
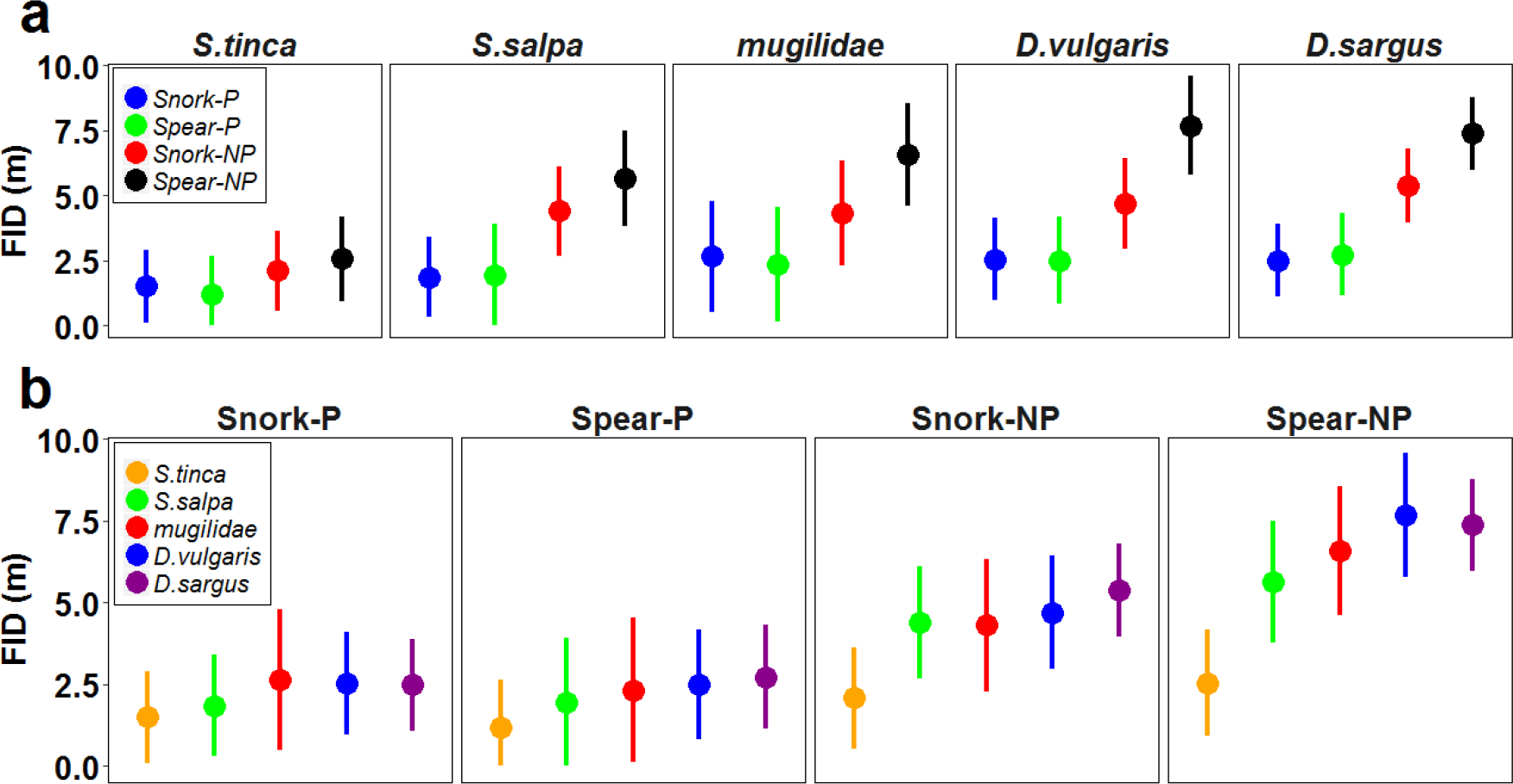
The results of the linear mixed model implemented for FID are reported to better visualize the information shown in Table 3. The estimation of FID are related to the significant three way interactions reported in Table 2 (Size × Treatment × Species). The mean estimation of FID adjusted for the average (22.5 cm) size effect are reported: (a) within the same species for each treatment; (b) within the same treatment for each species. In both cases, species and treatments are listed according to an increasing gradient of spearfishing pressure and threat, respectively. Vertical lines indicate the 95% confidence interval.

The presence of a snorkeler inside the MPA triggered a larger FID in *D. sargus* than in *S. tinca* (Table 3, Figs. 2, 3b). Inside the MPA, in presence of a spearfisher, the mean FIDs were significantly larger in *D. sargus* and *D. vulgaris* than in *S. tinca* (Table 3, Figs. 2, 3b). Outside MPA, in presence of a snorkeler, the FID was significantly shorter for *S. tinca* compared to the other four species/species groups (Table 3, Figs. 2, 3b). Finally, outside the MPA in the presence of a spearfisher, *D. sargus* and *D. vulgaris* showed significantly elevated FID, while *mugilidae* and *S. salpa* revealed intermediate FID, and *S. tinca* the shortest FIE (Table 3, Figs. 2, 3b).

The positive linear correlation between size and FID indicated that in almost all circumstances larger fish were more timid (Table 2, Fig. 2). The comparison within species indicated that the size effect increased outside the MPA only in two taxa (Table 3, Figs. 2, 4a): *mugilidae* (stronger effect of size in the presence of a snorkeler), and *D sargus* (stronger effect of size on FID both in the presence of a snorkeler and a spearfisher). The comparison within treatment also indicated that the size effect was significantly more pronounced in *D. sargus* than in *S. tinca* in the presence of a spearfisher inside the MPA and in the presence of a snorkeler outside the MPA (Table 3, Figs. 2, 4b). In the presence of a spearfisher outside the MPA, the effect of size was highly variable among species: stronger effects were recorded in the most targeted *D. sargus* and *D. vulgaris*, intermediate effects in the intermediately targeted *mugilidae* and *S. salpa* and the weakest effect on FID in the least targeted *S. tinca* (Table 3, Figs. 2, 4b).

**Figure 4.**
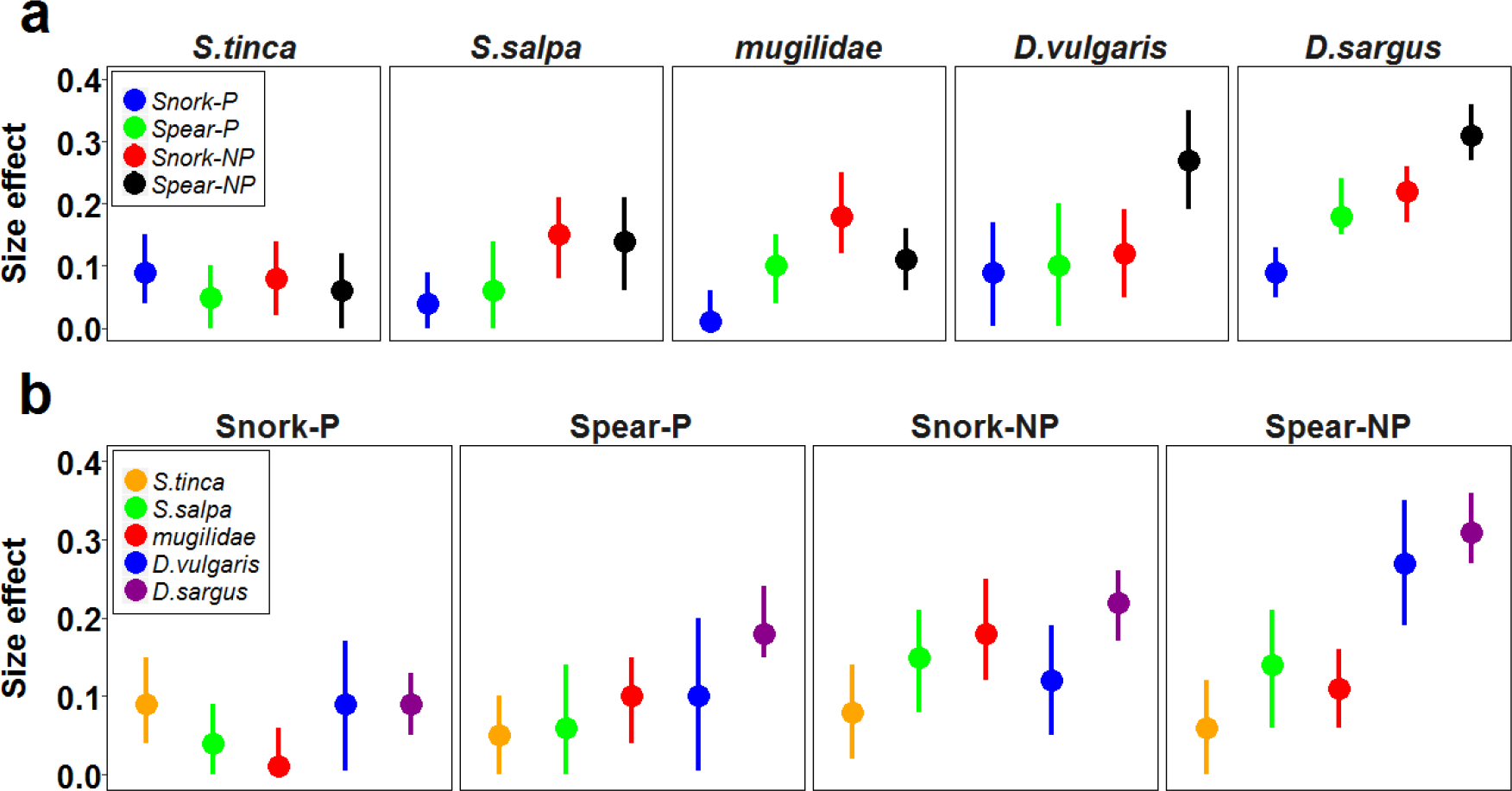
The results of the linear mixed model implemented for FID are reported to better visualize the information present in Table 3. The estimation of the FID are related to the size effect of three way interactions reported in Table 2 (Size × Treatment × Species). The estimation of the linear correlation between size and FID (slopes) are reported: (a) within the same species for each treatment; (b) within the same treatment for each species. In both cases, species and treatment are listed according to an increasing gradient of spearfishing pressure and threat, respectively. Vertical lines indicate the 95% confidence interval.

## DISCUSSION

We found that spearfishing promotes a timidity syndrome in exploited fishes and we confirmed all the three predictions we initially formulated. We found the risk perception of fishes changed with the appearance of the diver (spearfisher *vs* snorkeler) in a way commensurate with the level of threat posed by each of these two types of divers. Such differences were most evident outside the MPA, where fishes have historically experienced more encounters with threatening spearfishers and where certain behavioral types and morphotypes were selectively harvested, supporting the hypothesis of a spearfishing-induced timidity syndrome (Arlinghaus et al., 2017, Arlinghaus et al., 2016). Our finding that the timidity response was more pronounced not only in the historically most targeted species (e.g., *D. sargus* compared to *S. tinca*) but also in the largest fish further underscored that spearfishing has promoted increased shyness in the most targeted species and sizes. The positive correlation between FID and individual size can most plausibly be explained by experiential effects, but might also carry an evolutionary component, similar to findings reported from a freshwater salmonid targeted by recreational anglers in Japan (Tsuboi et al., 2016).

### Effect of spatial protection status against harvesting on FID

Protection of aquatic areas against exploitation has well known effects on fish behavior and typically has resulted in a decrease of fish timidity in a range of case studies ranging from spearfishing (Gotanda et al., 2009, Januchowski-Hartley et al., 2012, Januchowski-Hartley et al., 2011, Tran et al., 2016) to angling (Alós et al., 2014, Alós et al., 2015, Colefax et al., 2016, Cooke et al., 2017, Twardek et al., 2017). Agreeing with our results, previous studies have also shown that this response is species-specific and mainly shown by the most intensively targeted species (e.g. Colefax et al., 2016, Alós et al., 2015). We found that outside the MPA, the mean FID value were always larger than 4 m (except for *S. tinca*), which constitutes the maximum shooting range of the most commonly used spearguns (personal observations: VS, LM, EA) and strongly reduces the effectiveness of underwater human predation and elevates fish survival and hence fitness in the harvested areas. This finding means that common metrics used to assess fish abundance, such as catch rates or visual census, might strongly bias the perception about true fish abundance and negatively affect fisher satisfaction (Alós et al., 2015, Arlinghaus et al., 2017). Our findings also underscore the proposition that fish wariness assessed either visually or by underwater cameras can be used as a sensitive indicator of changes in fishing pressure (Goetze et al., 2017). Our findings in spearfishing can probably be extended to other recreational fishing gears (Tsuboi et al., 2016, Alós et al., 2015) and may well apply to many commercial gears as well (Arlinghaus et al., 2017).

### Risk recognition of human-induced predation threat

Changes in the appearance of the observer (spearfisher *vs* snorkeler) were expected to generate different responses among the targeted species. Our results demonstrate that fishes were able to discriminate between different level of threats (spearfisher vs snorkeler), and this ability varied among species according to their historical exposure to harvesting, body size and level of protection against harvesting. Fishes are therefore able to differentially sense human-induced predation risk and fine tune their behavioral response (Frid and Dill, 2002). In the case of spearfishing, the optimal FID navigates the fundamental trade-off among energy acquisition (e.g., time spent foraging) and the risk of being captured or attacked by natural predators (Ydenberg and Dill, 1986), which has strong repercussions for individual fitness (Frid and Dill, 2002). Although fishes are generally known to accomplish this trade-off when being exposed to natural predators (e.g. Werner and Hall, 1974), our work shows that the same is also valid for underwater human predators. For example, among the studied species, the historically most exploited ones, *D. sargus* and *D. vulgaris* (Lloret et al., 2008, Rocklin et al., 2011), showed a steep increase of FID along the gradient of human-induced predation risk. The recognition of spearfishers as a potential predator adds important information to the generally small and partly contradictory body of literature on this topic. For example, the presence of a speargun was not associated with a specific response in exploited parrotfishes in a previous study (Januchowski-Hartley et al., 2012), whilst the unexploited lined bristletooth (*Ctenochaetus striatus*) was found to be able to discriminate between a snorkeler with or without a speargun (Tran et al., 2016). Our results support the hypothesis that the response to spearfishers is species-specific and probably harvest-dependent, and that the most heavily exploited species and fish sizes within a species are able to use visual cues to assess human-predation risk and properly allocate resources to avoid the risk to being harmed or captured, similar to the case in natural predation (Kelley and Magurran, 2003, McCormick and Manassa, 2008). Also, a sizable literature on catch-and-release recreational angling has shown that fishes are able to develop rapid hook avoidance (Klefoth et al., 2013) and that previous catch-and-release results in behavioral alteration and the seeking of refugees (Klefoth et al., 2008).

### The ability of different sized fishes to respond to human-induced predation risk

The positive relationship between individual size and FID was a common pattern in all the five studied species. This result might be explained by a lower risk-taking tendency associated with the higher reproductive value of large individuals (Gotanda et al., 2009) or simply be a natural outcome of learning processes through development. The finding can also be explained by fisheries-induced evolution of increased timidity that is most expressed in the most vulnerable morphological or ontogenetic stage. Our field methods were not intended to differentiate among behavioral plasticity and evolutionary responses. Nevertheless, size effects were significantly stronger in highly exploited fish species (specifically *D. sargus*), and this could support a plastic hypothesis. Therefore, the observed changes in FID should be mainly caused by previous experience with spearfishers across ontogeny, even if evolutionary explanations cannot be ruled out. In the MPAs the size effect in *D. sargus* was stronger in presence of a spearfisher than when confronting a non-threatening snorkeler. This may be explained by experiences acquired beyond the boundaries of protected areas, because some *D. sargus* individuals are known to visit areas outside the MPAs (e.g. Di Lorenzo et al., 2014) where they may experience spearfishing and learn. Otherwise, larger individuals could have experienced spearfishing predation risk inside the MPA due to poaching activities. Beside the relative contribution of these two not mutually exclusive explanations, such sharp differences in the size effect within and outside MPA confirm the fish wariness as a suitable indicator of human-induced predation risk in selected species (Goetze et al., 2017).

### Mechanisms underlying the FID responses

The timidity syndrome caused by exploitation has been explained by two main mechanisms (Arlinghaus et al., 2017): the first is related to a genetic basis for those behaviors that contribute to fishing vulnerability, as demonstrated in recreational angling (Klefoth et al., 2013, Philipp et al., 2009). This means that selective forces induced by human predation are operating on genetic components of those traits controlling FID, leading to fisheries-induced evolution of behavior (Heino et al., 2015, Uusi-Heikkilä et al., 2008). The second mechanism relates to a purely plastic behavioral response of fish that is mediated by learning processes to avoid exposure to fishing. This can be achieved by both individual (private, e.g., catch-and-release or spearfishing-induced injury) and social experiences (e.g., seeing conspecifics being injured or captured). It is highly likely that, similar to the case in life-history traits, behavioral plasticity overrules evolutionary effects, but previous common-garden-experiments using both laboratory models as well as wild-living populations raised in laboratories have shown that size-selective harvesting can indeed evolutionarily lead to timidity (Uusi-Heikkilä et al., 2015, Tsuboi et al., 2016).

On different timescales, both plastic and evolutionary responses may contribute to the modulation of FID. However, considering the broad connectivity of marine systems and the often-observed absence of genetic structuring for many marine fish populations (see Sahyoun et al., 2016, Pujolar et al., 2013), the observed increase of FID outside the MPA seems more likely explained by learning processes than by genetic causes. A stronger role of learning mechanisms is also supported by the fact that periodically harvested fishing closures resulted in a decrease of FID (Januchowski-Hartley et al., 2014). Nevertheless, the hypothesis of an ongoing spearfishing-induced selection of FID cannot be excluded and should be tested by common garden experiments or by reciprocal transplants in the wild (Diaz Pauli and Heino, 2014).

## CONCLUSIONS

Our study provides a comprehensive assessment of how spearfishing modulates fish anti-human-predator behavior in the wild commensurate with the level and intensity of harvesting pressure directed at given species, sizes and sites. The minimum distance to which a fish can be approached by a human predator can be related conceptually to the safe operating space concept developed by Rockstrom et al. (2009). We suggest that spearfishing systematically increases the safe operating distance of fishes, which benefits individual fitness and conservation and harms spearfisher`s catch rates. The behavior adopted by the observer in this study (chasing the fishes from the surface) constitutes an active harvesting technique where the ultimate decision to catch a fish is taken by the spearfisher. This demonstrates that a timidity syndrome is also possible by means of active fishing gears and not only by passive fishing gear types as previously argued (Diaz Pauli and Sih, 2017, Arlinghaus et al., 2017). If spearfishers want to maintain high catch rates, either implementation of rotating closures, careful constraints on local spearfishing pressure and implementation of a network of marine protected areas with spill-over of naïve fishes to unprotected sites (Januchowski-Hartley et al., 2013) can be recommended. These means should recover boldness of previously exploited fishes and maintain catch rates high. However, if spearfishing systems remain unmanaged or poorly managed, constant erosion of catchability is likely to occur, which reduces the index-value of spearfishing data to index abundance and in extreme cases may also harm non-threatening snorkelers by reducing visual contacts with fish. Such changes could fuel competition among spearfishers and lead to social and political conflict.

## Acknowledgments

We are grateful to the management body of Bonifacio Straits Natural Reserve, and warmly acknowledge the support provided by Michel Culioli. We also tank the MPAs of Cerbère-Banyuls and Medes Islands for their availability and logistical support. VS is supported by a Leibniz-DAAD postdoctoral research fellowship (# 91632699).

## Authors’ contribution

EA conceived and designed the study with the contribution of LM; LM acquired data; VS, LB, BW supported during acquisition of data; VS and RA statistically analysed the data, VS, EA and RA interpreted the data, and VS led the writing of the manuscript with inputs by all other co-authors. All authors gave final approval for publication.

## Data Accessibility

Data will be available on Dryad (http://datadryad.org/) upon acceptance of the paper.

